# Yeast-based production platform for potent and stable heavy chain-only antibodies

**DOI:** 10.1101/2024.03.04.580093

**Authors:** Chiara Lonigro, Hannah Eeckhaut, Royan Alipour Symakani, Kenny Roose, Bert Schepens, Koen Sedeyn, Anne-Sophie De Smet, Jackeline Cecilia Zavala Marchan, Pieter Vanhaverbeke, Sandrine Vanmarcke, Katrien Claes, Sieglinde De Cae, Hans Demol, Simon Devos, Daria Fijalkowska, Wim Nerinckx, Iebe Rossey, Wannes Weyts, Rana Abdelnabi, Dirk Jochmans, Johan Neyts, Xavier Saelens, Loes van Schie, Nico Callewaert

**Author notes:** Correspondence should be addressed to N.C. These authors contributed equally to this manuscript (Chiara Lonigro, Hannah Eeckhaut).

## Abstract

Monoclonal antibodies are the leading drug of the biopharmaceutical market because of their high specificity and tolerability, but the current CHO-based manufacturing platform remains expensive and time-consuming leading to limited accessibility, especially in the case of diseases with high incidence and pandemics. Therefore, there is an urgent need for an alternative production system.

In this study, we present a rapid and cost-effective microbial platform for heavy chain-only antibodies (VHH-Fc) in the methylotrophic yeast *Komagataella phaffii* (aka *Pichia pastoris*). We demonstrate the potential of this platform using a simplified single-gene VHH-Fc fusion construct instead of the conventional monoclonal antibody format, as this is more easily expressed in *Pichia pastoris*. We demonstrate that the *Pichia*-produced VHH-Fc fusion construct is stable and that a *Pichia*-produced VHH-Fc directed against the SARS-CoV-2 spike has potent SARS-CoV-2 neutralizing activity *in vitro* and *in vivo*. We expect that this platform will pave the way towards faster and cheaper development and production of broadly neutralizing single-chain antibodies in yeast.

## Main text

Antibodies represent a well-studied class of therapeutics for the treatment and prevention of infectious diseases. Early administration of recombinant neutralizing monoclonal antibodies (mAbs) is able to avert severe disease. Unlike active immunization, passive immunization with monoclonal antibodies offers immediate protection and therefore can be administered prophylactically to achieve ring immunization to respond to an outbreak of a new infectious disease. In this context, rapid microbial manufacturing at lowered cost of goods would be an asset^1^.

Yeast-based manufacturing of mAbs is attractive because of the fast, low-cost and scalable fermentation processes^2^. Furthermore, the first yeast-produced mAb has been recently market approved, opening up a regulatory approval track for this production system^3^. Unfortunately, secretion of conventional IgG antibodies in yeast is not trivial^4^. Successful expression of full-length antibodies in *Pichia pastoris* has been reported for only a few mAbs^5,6^, yet most of the data refer to the anti-Her2 antibody trastuzumab (Herceptin), which is known in the field to be an easy antibody to express.

Therefore, we decided to focus our attention on a simplified single-gene VHH-Fc fusion construct (Figure 1 A). We previously identified a VHH that can neutralize both SARS-CoV-1 and SARS-CoV-2, called VHH72^7^, and detailed its conversion into the first-reported potent anti-SARS-CoV-2 mAb XVR011^8^. In the latter study, we used the *Pichia pastoris* expression system to rapidly screen affinity-enhanced VHH-Fc mutants, directly in the drug-relevant Fc fusion format, such that only mutants with an increased affinity/avidity in this bivalent format were identified and rapidly moved forward in the drug development campaign. The data showed that the expression of VHH-Fc in *Pichia pastoris* is robust and flexible, which motivated us to explore it further as a valuable alternative to the current CHO-based manufacturing platform by further improving engineered Fc variants for *Pichia*.

**Figure 1.**
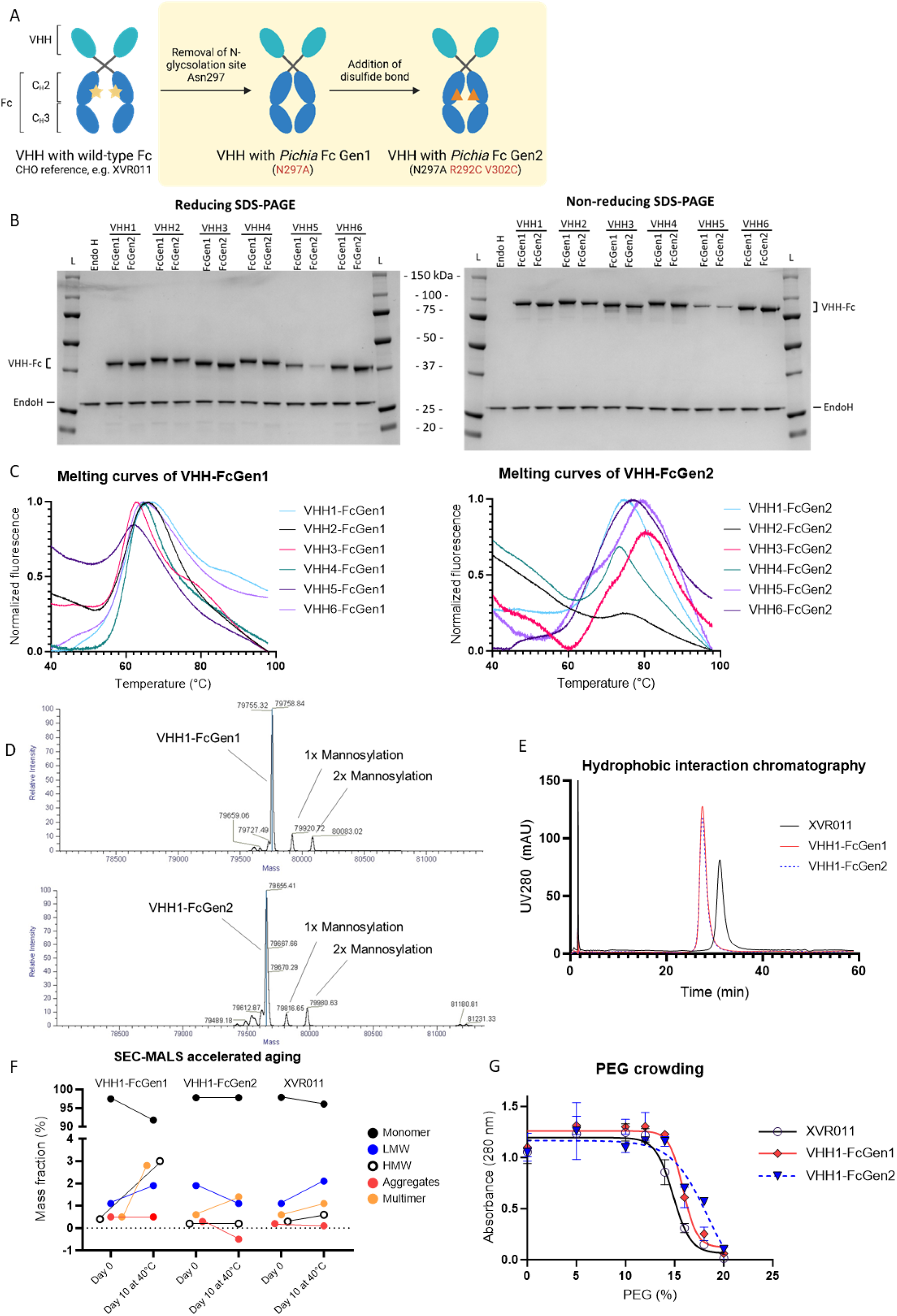
Manufacturability and biophysical properties of Fc-engineered VHH-Fc molecules produced in *Pichia pastoris*. A. Schematic overview and nomenclature of the molecules used in this study. Glycosylation is indicated by a yellow star, and an additional disulfide bond with an orange triangle. B. Reducing and non-reducing SDS-PAGE of multiple VHH-Fcs produced by *Pichia pastoris* strain OPENPichia with P_GAP_ promotor and purified via protein A and size exclusion chromatography. 0.5 μg of purified protein was loaded. EndoH (0.25 μg) was used as a loading control. Ladder (L) is Precision Plus Protein™ All Blue Prestained Protein Standards. C. Thermostability of multiple VHH-Fcs by SYPRO Orange thermoshift assay. Fluorescence was normalized via GraphPad Prism. D. LC-MS spectra of intact VHH-Fcs produced by *Pichia pastoris* and purified via protein A and size exclusion chromatography. The raw spectra were deconvoluted with the ReSpect Algorithm in BioPharma Finder software, followed by manual annotation. E. Protein hydrophobicity was evaluated by hydrophobic interaction chromatography (HIC) on a Dionex ProPac HIC-10 column column. The separation was monitored by measuring Abs at 280 nm. F. SEC-MALS analysis of purified VHH1-Fc samples before and after 10 days at 40 °C. The peak corresponding to the molar weight of an assembled VHH-Fc protein was indicated as ‘monomer peak’. Peak quantitation (%) is based on the refraction signal. Qualitative analysis of the monomer peak was performed on the 200 μl peak elution fraction. Values reported are from a single stress tested sample. HMW = high molecular weight species; LMW = low molecular weight species. G. Aggregation tendency was evaluated by incubating the protein with increasing concentrations of PEG3350 and measuring the concentration of soluble protein by measuring Abs at 280 nm (± SD, n=3).

We strategically engineered the small and single chain VHH-Fc antibody format, a genetic fusion of the highly stable variable heavy domain of llama-derived heavy-chain only antibodies (VHHs), to the human immunoglobulin G1 (hIgG1) Fc. We adapted the hIgG1 sequence to better match yeast expression (Figure 1 A, Supplementary Figure S1). To start, the first five amino acids of the hinge (EPKSC) were truncated to remove the unpaired cysteine that is usually involved in a disulfide bond with the light chain in a conventional full-length antibody. Secondly, the Fc C-terminal lysine was kept, since, in contrast to CHO produced IgGs, proteolytic processing of this residue does not occur in *Pichia*. Finally, differences in the N-glycosylation pathway between yeast and humans have so far hampered its use for production of Fc containing molecules for human use. The yeast N-glycosylation machinery produces glycoproteins modified with yeast-specific high-mannose oligosaccharide N-glycans rather than mammalian complex-type N-glycans, which leads to rapid clearance after injection and may be immunogenic. Mutation of Asn297 (EU numbering^9^) in the Fc-C_H_2 domain results in an aglycosylated Fc, which is safe for medical use^3^. As the aglycosylated Fc does not bind FcγRs and C1q, there was no longer a need to introduce mutations to abolish effector function. We chose the N297A substitution, since the same mutation is present in the Fc of Vyepti (eptinezumab), the first aglycosylated and *Pichia-* produced mAb approved on the market^3^. We named the construct resulting from this Fc engineering campaign “(*Pichia*) FcGen1” (Figure 1 A).

Previous studies reported that the absence of the sugar chain that otherwise occupies the space between the paired Fc-C_H_2 domains reduces the stability of the VHH-Fc molecule as compared to the glycosylated IgG^10,11^. It has been shown that the introduction of an additional disulfide bond in an isolated aglycosylated C_H_2 domain restores or even increases the thermal stability^10^. Therefore, as an attempt to compensate for the stability loss, an extra disulfide bond was introduced in the C_H_2 domain of our VHH-Fc construct by introducing substitutions R292C and V302C^11^ (Figure 1 A and Supplementary Figure 1). We named the resulting construct “(*Pichia*) FcGen2” (Figure 1 A).

Subsequently, the VHH-Fcs were produced in *Pichia* and thoroughly characterized. To broaden our scope, we started from a large and diverse set of VHHs to test in parallel with both engineered Fc constructs (named “VHH1 to 6”). First, we show that the addition of a disulfide bond to the Fc has no apparent influence on the secretability of the molecules, as evidenced from a random clone screening on SDS-PAGE (Supplementary Figure S2). We also show that the Fc-engineered molecules can be purified using protein A chromatography and subsequent size-exclusion chromatography with adequate yields (Figure 1 B, Supplementary Figures S3). We confirmed the molecular identity of the purified products using intact mass spectrometry (MS) (Figure 1 D). Intact MS also revealed O-mannosylation, as well as the conversion of the N-terminal Gln to Pyro-Glu (the latter only in the case of VHH6).

Additionally, we characterized the molecules for their thermal stability, hydrophobicity, and aggregation tendency, as these are critical attributes in the development of a candidate biologic. As expected, the aglycosylated VHH-FcGen1 showed a reduced thermal stability in a thermoshift assay, as compared to the molecules with the additional disulfide bond as present in the FcGen2 construct (Figure 1 C). The addition of the disulfide bond in the Fc region did not impact the hydrophobicity (Figure 1 E). Of note, both *Pichia*-produced Fc-engineered constructs were less hydrophobic than reference CHO-produced XVR011, possibly because of the more compact conformation due to the absence of the N-glycans in the C_H_2 cavity (Figure 1 E). Additionally, the molecule with the stabilized aglycosylated Fc (FcGen2) showed a lower tendency of forming soluble and/or insoluble aggregates in accelerated aging experiments (10 days at 40°C), which is either equal or better as compared to CHO-produced N-glycosylated XVR011 (Figure 1 F). The *Pichia*-produced VHH-Fcs also showed a reduced hydrophobicity and aggregation tendency in the presence of high concentrations of PEG3350 (Figure 1 G). Overall, the *Pichia*-produced stabilized aglycosylated VHH-FcGen2 showed desirable physicochemical properties.

We decided to continue with the anti-Covid (VHH1 and VHH2) Fc-engineered VHH-Fc molecules for *in vitro* potency testing. Neither the binding of the different *Pichia*-produced VHH-Fc variants to SARS-CoV-2 RBD and spike, or the *in vitro* neutralization potency was affected by the Fc engineering (Figure 2 A-C). Interestingly, compared to XVR011, the *Pichia*-VHH-Fc constructs showed higher affinity for immobilized SARS-CoV-2 RBD and spike protein on ELISA, a slightly higher potency in VSV SARS-CoV-2 spike pseudotyped neutralization assay, and a similar potency in neutralizing of authentic SARS-CoV-2 virus *in vitro* and reducing plaque formation (Figure 2 D). Additionally, we show that the epitope of the *Pichia* produced VHH2-FcGen2 is very similar to the epitope of CHO-produced XVR011 based on deep mutational scanning^12^ (Figure 2 E).

**Figure 2.**
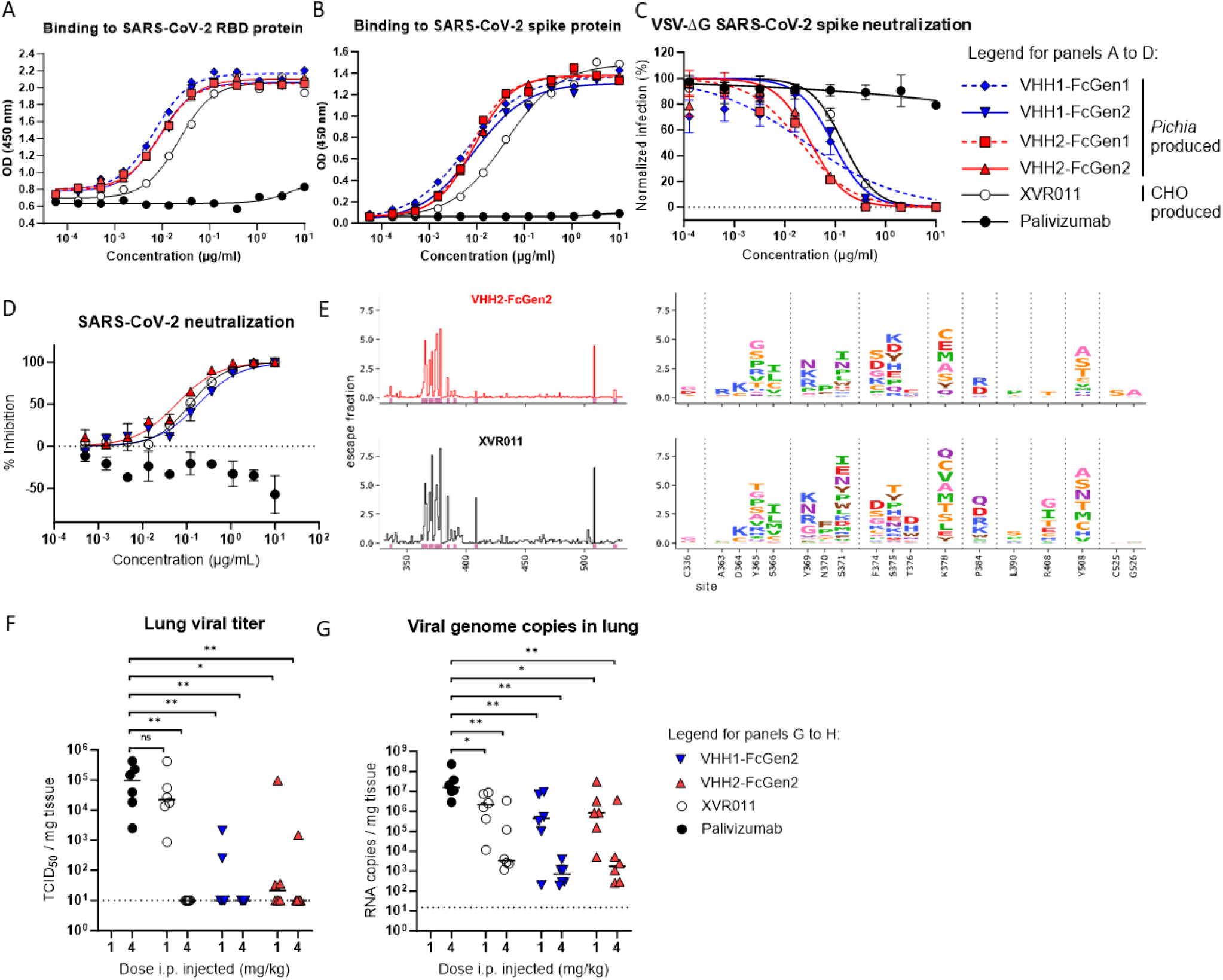
Potency of *Pichia pastoris*-produced Fc-engineered SARS-CoV-2 neutralizing VHH-Fc molecules. A. Binding of the indicated constructs to SARS-CoV-2 RBD determined by ELISA (data points are the mean; n=3). B. Binding of the indicated constructs to SARS-CoV-2 spike determined by ELISA (data points are the mean; n=3). C. Neutralization of the indicated constructs of SARS-CoV-2 S pseudotyped VSV. The graphs show the mean (± SD, n=4) GFP fluorescence normalized to the mean GFP fluorescence of noninfected and infected PBS-treated cells. D. SARS-CoV-2 BetaCov/Belgium/GHB-03021/2020 plaque reduction neutralization assay by the indicated constructs (± SD, n=2). Palivizumab is an RSV-neutralizing mAb. E. Deep mutational scanning profiles against yeast surface-displayed SARS-CoV-2 RBD libraries. The escape fractions are represented in logo plots, with letter height corresponding to the level of escape (n = 2). F. – G. Hamsters received 1 or 4 mg/kg of XVR011, VHH1-FcGen2, VHH2-FcGen2, or 4 mg/kg of Palivizumab by intraperitoneal injection 24 hours after intranasal administration of 2×10^6^ TCID_50_ SARS-CoV-2 BetaCov/Belgium/GHB-03021/2020. Lung samples were collected at day 4 post-infection from euthanized hamsters and viral RNA and infectious virus were quantified by RT-qPCR and end-point virus titration, respectively. Data were analyzed with the Kruskal-Wallis test and Dunn’s multiple comparison test (*P<0.05; **P<0.01). Horizontal bars indicate median. Dotted horizontal lines indicate lower limit of detection.

Next, we tested the *in vivo* anti-SARS-CoV-2 potency of the constructs in hamsters. Doses of 1 and 4 mg/kg of VHH1-FcGen2, VHH2-FcGen2 or XVR011 were administered intraperitoneally 24 hours after challenge. Control animals received 4 mg/kg of the respiratory syncytial virus targeting palivizumab (Synagis). No infectious virus was detectable in lung homogenates from any of the 4 mg/kg VHH-Fc treated hamsters except for one outlier, which was later determined to be caused by a misinjection (Figure 2 F-G). Surprisingly, the *Pichia* constructs significantly reduced the titer of infectious virus at the lower dose of 1 mg/kg as well, whereas XVR011 did not. The two-fold increase in *in vitro* potency of the VHH2-FcGen2 over the VHH1-FcGen2 construct did not translate into a higher *in vivo* potency. However, both *Pichia*-made constructs showed higher reduction of infectious virus levels in the lungs at 1 mg/kg, when compared to CHO-made lab grade XVR011, which did not differ from the negative control-treated hamsters at this low dosage. This might indicate a more favorable lung biodistribution of *Pichia*-produced VHH1-FcGen2 and VHH2-FcGen2.

In conclusion, the development and production of antibody-like biopharmaceuticals in yeast offers a unique opportunity to speed up the drug discovery process and reduce the manufacturing costs of goods, and to drive towards a broader commercial utilization of these drugs. This solidifying work establishes a promising track for the design of homogenous, highly stable, and potent heavy chain-only antibodies that can be produced in *Pichia pastoris*.

## Materials and methods

### Strains, media and reagents

*Escherichia coli* (*E. coli*) MC1061 was used for standard molecular biology manipulations. For plasmid propagation, *E. coli* were cultured in LB broth (0.5% yeast extract, 1% tryptone, and 0.5% NaCl) supplemented with 25 μg/mL chloramphenicol (MP Biomedicals), 50 μg/mL carbenicillin (Duchefa Biochemie) or 50 μg/mL Zeocin (Life Technologies). *P. pastoris* NCYC 2543 strain was provided by the National Collection of Yeast Culture. *P. pastoris* strain OPENPichia has been described before^13^. Yeast cultures were grown in liquid YPD (1% yeast extract, 2% peptone, 2% D-glucose) or on solid YPD-agar (1% yeast extract, 2% peptone, 2% D-glucose, 2% agar) at pH 7.5 and selected with 100 μg/mL Zeocin. For protein expression, cultures were grown in a shaking incubator (28°C 225 rpm) in BMGY (same composition but with 1% glycerol replacing the 2% D-glucose) or BMMY (same composition but with 1% methanol replacing the 2% D-glucose).

### Generation of expression plasmids

Most of the expression vectors for the VHH-Fc variants were generated using an adapted version of the Yeast Modular Cloning toolkit based on Golden Gate assembly^13,14^. Briefly, coding sequences for the VHH and hIgG-Fc variants, were codon optimized for expression in *P. pastoris* and ordered as gBlocks at IDT. Each coding sequence was flanked by unique part-specific upstream and downstream BsaI-generated overhangs. The gBlocks were inserted in a universal entry vector via BsmBI assembly which resulted in different “part” plasmids (entry vectors), containing chloramphenicol resistance cassette. Part plasmids were assembled to form expression plasmids (pX-VHH-Fc) via a Golden Gate BsaI assembly.

Each expression plasmid consists of the assembly of 9 parts: P1_ConLS, P2_pGAP (or P2_pAOX1), P3-Ost1-VHH (or P3-Kar2-VHH), P4a-hIgG-Fc, P4b_AOX1tt, P5_ConR1, P6-7 Lox71-Zeo or P6-7 Stuffer, P8 AmpR-ColE1-Lox66 or P8 Ori-Zeo. Selection of correctly assembled expression plasmids was performed in LB supplemented with either 50 μg/mL carbenicillin and 50 μg/mL Zeocin. All the entry vectors and expression plasmids were sequence verified. Point mutations N297A, R292C and V302C were introduced by site directed mutagenesis using QuikChange Lightning Site-Directed Mutagenesis Kit (Agilent) according to the manufacturer’s protocol.

### Generation of *Pichia pastoris* expression strains

Prior to transformation, all expression constructs were linearized in the promoter region (AvrII for P_GAP_ and PmeI for P_AOX1_) to facilitate single homologous recombination and targeted integration in the *P. pastoris* genome. Transformations were performed using the lithium acetate electroporation protocol as described by Wu and Letchworth^15^. Briefly, a culture of *P. pastoris* was grown overnight in YPD to obtain an OD_600_ of 1.5. The cells were harvested and were incubated in 100 mM LiAc, 10 mM DTT, 0.6 M sorbitol and 10 mM Tris-HCl, pH 7.5. The cells were thoroughly washed with 1 M ice-cold sorbitol. Cells were transformed by electroporation (1.5 kV, 25 μF, 200 Ω) with 75-1000 ng of linearized vector. Positive transformants were selected on YPD-agar supplemented with 100 μg/mL of Zeocin.

### Protein expression and purification

For protein expression, an overnight culture of *P. pastoris* transformed with the expression plasmid was diluted in BMGY to 0.1 OD_600_ in baffled shake flasks. For methanol-based expression with P_AOX1,_ after 48h of growth in glycerol, expression was induced by switching to medium containing 1% MeOH. To maintain induction, cultures were spiked with 1% MeOH every 8-12h. For expression with the constitutive *GAP* promoter, expression was performed in BMGY for 50-60h, at 28 °C, 250 rpm. Medium was collected by centrifugation at 1,500 g, 4 °C for 10 minutes and filtered over a 0.22 μm bottle top filter (Millipore) before loading on a HiTrap MabSelect SuRe 5 ml column (GE Healthcare), equilibrated with McIlvaine buffer pH 7.2 (174 mM Na_2_HPO_4_, 13mM citric acid). The column was eluted with McIlvaine buffer pH 3 (40 mM Na_2_HPO_4_, 79mM citric acid). Collected fractions were neutralized to pH 6.5 with 0.4M Na_3_PO_4_. The elution fractions were analyzed on SDS-PAGE. Target protein-containing fractions were pooled and finally, the protein was injected on a HiLoad 16/600 Superdex 200 pg column (GE-Healthcare) or Cytiva Superdex 200 10/300 GL column (GE-Healthcare), and eluted with PBS. The obtained fractions were analyzed by SDS-PAGE and the fractions containing monomeric VHH-Fc were pooled together. Protein concentration was measured by 280 nm absorbance versus a buffer blank and concentrated with Amicon 30 kDa MWCO spin columns. Purified protein was snap frozen in liquid nitrogen and stored at -80 °C.

### Mass spectrometry analysis of proteins

Intact VHH-Fc proteins were separated on an Ultimate 3000 HPLC system (Thermo Fisher Scientific) online connected to an LTQ Orbitrap Elite mass spectrometer (Thermo Fischer Scientific). Briefly, approximately 2 μg of protein was injected on a Zorbax Poroshell 300SB-C8 column (5 μm, 300Å, 1x75mm IDxL; Agilent Technologies) and separated using a 15 min gradient from 5% to 80% solvent B at a flow rate of 100 μl/min (solvent A: 0.1% formic acid and 0.05% trifluoroacetic acid in water; solvent B: 0.1% formic acid and 0.05% trifluoroacetic acid in acetonitrile). The column temperature was maintained at 60°C. Eluting proteins were directly sprayed in the mass spectrometer with an ESI source using the following parameters: spray voltage of 4.2 kV, surface-induced dissociation of 30 V, capillary temperature of 325 °C, capillary voltage of 35 V and a sheath gas flow rate of 7 (arbitrary units). The mass spectrometer was operated in MS1 mode using the orbitrap analyzer at a resolution of 30,000 (at m/z 400) and a mass range of 600-4000 m/z, in profile mode. The resulting MS spectra were deconvoluted with the BioPharma FinderTM 5.2 software (Thermo Fischer Scientific) using the ReSpect deconvolution algorithm (isotopically unresolved spectra). The deconvoluted spectra were manually annotated.

### Thermal stability

To evaluate thermal stability of VHH-Fc variants, differential scanning fluorimetry (a thermofluor assay) was performed. Briefly, a final concentration of 0.1 μg/ml of VHH-Fc in PBS was mixed with 10X SYPRO Orange dye (Life Technologies). Dye binding to molten globule unfolding protein was measured over a 0.01 °C/s temperature gradient from 20 °C to 98 °C in a Roche LightCycler 480 qPCR machine.

### Hydrophobic interaction chromatography (HIC)

Apparent hydrophobicity was assessed using a HIC assay employing a Dionex ProPac HIC-10 column, 100 mm×4.6 mm (Thermo Fisher 063655), containing a stationary phase consisting of a mixed population of ethyl and amide functional groups bonded to silica. All separations were carried out on an Agilent 1200 HPLC equipped with a fluorescence detector. The column temperature was maintained at 20 °C throughout the run and the flow rate was 0.8 ml/min. The mobile phases used for the HIC method were (A) 0.8 M ammonium sulfate and 50 mM phosphate pH 7.4, and (B) 50 mM phosphate pH 7.4. Following a 2 min hold at 0% B, the column was loaded with 15 μl of sample at 2 mg/ml, and bound protein was eluted using a linear gradient from 0 to 100% B in 45 min and the column was washed with 100% B for 2 min and re-equilibrated in 0% B for 10 min prior to the next sample. The separation was monitored by absorbance at 280 nm.

### SEC MALS

To determine the molecular mass and aggregation behavior of VHH-Fc variants, the protein was analyzed by size exclusion chromatography multi-angle laser light scattering (SEC-MALS). For each analysis, 100 μl sample filtered through 0.1μm Ultrafree-MC centrifugal filters (Merck) was injected onto a Superdex200 10/300 GL Increase SEC column (GE Healthcare) equilibrated with sample buffer, coupled to an online UV detector (Shimadzu), a mini DAWN TREOS (Wyatt) multi-angle laser light scattering detector and an Optilab T-rEX refractometer (Wyatt) at 298 K. The refractive index (RI) increment value (dn/dc value) at 298 K and 658 nm was calculated using SEDFIT v16.175 ^16^ and used for the determination of the protein concentration and molecular mass.

### Polyethylene glycol (PEG) aggregation assay

A 40% stock PEG 3350 (Merck, 202444) solutions (w/v) were prepared in PBS pH 7.4. To minimize non-equilibrium precipitation, sample preparation consisted of mixing protein and PEG solutions at a 1:1 volume ratio. 35 μL of a 2 mg/ml sample solution was added to the PEG stock solutions resulting in a 1 mg/ml test concentration. Samples were incubated at 20 °C for 24 h. The sample plate was subsequently centrifuged at 4000 x g for 1 h at 20 °C. 50 μL of supernatant was dispensed into a UV-Star, half area, 96 well, μClear, microplate (Greiner, 675801). Protein concentrations were determined by UV spectrophotometry at 280 nm using a FLUOstar Omega multi-detection microplate reader (BMGLABTECH). The resulting values were plotted using GraphPad Prism 7.04 and the PEG midpoint score was derived from the midpoint of the sigmoidal dose-response (variable slope) fit.

### ELISA

Wells of microtiter plates (type II, F96 Maxisorp, Nunc) were coated overnight at 4 °C with 100 ng of recombinant SARS-CoV-2 S-2P protein (with foldon), mouse Fc-tagged SARS-CoV-2 RBD (Sinobiological) or BSA. The coated plates were blocked with 5% milk powder in PBS. Dilution series of the VHHs were added to the wells. Binding was detected by incubating the plates sequentially with HRP-linked rabbit anti-human IgG (A8792, Sigma). After washing, 50 μL of TMB substrate (Tetramethylbenzidine, BD OptETA) was added to the plates and the reaction was stopped by addition of 50 μL of 1 M H_2_SO_4_. The absorbance at 450 nm was measured with an iMark Microplate Absorbance Reader (Bio Rad). Curve fitting was performed using nonlinear regression (Graphpad Prism 9.1).

### VSV pseudovirus neutralization

To generate replication-deficient VSV pseudotyped viruses, HEK293T cells, that were transfected with an expression vector encoding the spike protein of SARS-CoV-1 S, SARS-CoV-2 S or the mutants thereof were inoculated with a replication deficient VSV vector containing eGFP and firefly luciferase expression cassette ^17,18^. After a 1 h incubation at 37 °C, the inoculum was removed, cells were washed with PBS and incubated in media supplemented with an anti-VSV G mAb (ATCC) for 16h. Pseudotyped particles were then harvested and clarified by centrifugation ^19^. VSV particles were subsequently PEG-precipitated form the cleared medium and resuspended in FluoroBirte DEMEM medium (Thermo Fisher Scientific) containing 5% fetal calf serum (FCS). For the VSV pseudotype neutralization experiments, pseudoviruses were incubated for 30 min at room temperature with different concentrations of antibodies. The incubated pseudoviruses were subsequently added to confluent monolayers of Vero E6 cells. Sixteen hours later, the cells were lysed using passive lysis buffer (Promega). The transduction efficiency was quantified by measuring the GFP fluorescence in the prepared cell lysates using a Tecan infinite 200 pro plate reader. GFP fluorescence was normalized using either the GFP fluorescence of non-infected cells and infected cells treated with PBS. The IC50 was calculated by non-linear regression curve fitting, log(inhibitor) versus normalized response using Graphpad Prism 9.1.

### Plaque reduction neutralization test (PRNT)

We used SARS-CoV-2 strain BetaCov/Belgium/GHB-03021/2020 (EPI ISL 407976; 2020-02-03) used from passage P6 grown on Vero E6 cells. Dose-dependent neutralization of distinct VHH-Fc constructs was assessed by mixing the VHH-Fc constructs at different concentrations (three-fold serial dilutions starting from a concentration of 10 μg/ml), with 100 PFU SARS-CoV-2 in DMEM supplemented with 2% fetal bovine serum (FBS) and incubating the mixture at 37 °C for 1h. VHH-Fc-virus complexes were then added to Vero E6 cell monolayers in 12-well plates and incubated at 37 °C for 1h. Subsequently, the inoculum mixture was replaced with 0.8% (w/v) methylcellulose in DMEM supplemented with 2% fetal bovine serum. After 3 days incubation at 37 °C, the overlays were removed, the cells were fixed with 3.7% PFA, and stained with 0.5% crystal violet. Half-maximum neutralization titers (PRNT50) were defined as the VHH-Fc concentration that resulted in a plaque reduction of 50% across 2 or 3 independent plates.

### Deep mutational scanning

Two independent deep mutational libraries of the SARS-CoV-2 receptor-binding domain (N331-T531) cloned in the pETcon vector were kindly gifted by Dr. Jesse Bloom^12^. Ten ng of each library was electroporated to electrocompetent *E. coli* TOP10 cells with the Gene Pulser electroporation system (BioRad) set at 200 Ω, a capacitance of 25 μF, a capacitance extension of 125 μF, and a voltage of 2.5 kV. Cells were transferred to SOC medium (2% tryptone, 0.5% yeast extract, 10 mM NaCl, 2.5 mM KCl, 10 mM MgCl_2_, and 20 mM glucose) and allowed to recover for 1h. Afterwards, a serial dilution was plated to assess transformation efficiency, and the remainder was plated on large low salt LB agar plates supplemented with ampicillin. After overnight incubation at 37 °C, colonies were scraped from the plates, inoculated in low salt LB medium supplemented with ampicillin and grown for 2 hours 30 min at 37 °C. Afterwards, the cultures were pelleted by centrifugation and washed once with sterile Milli-Q water. Plasmid was extracted via the QIAfilter plasmid Giga prep kit (Qiagen) according to the manufacturer’s instructions.

Ten μg of every extract was transformed to the *S. cerevisae* EBY100 strain following the large-scale protocol by Gietz and Schiestl^20^, and selected in liquid SD-trp – ura drop-out medium for 16h. Afterwards, the cultures were back-diluted at 1 OD_600_/mL in fresh SD –trp –ura drop-out medium for an additional 9h passage. A serial dilution was plated to assess transformation efficiency, and the remainder of the cultures were snap frozen at 1e8 cells per aliquot and stored at -80 °C for further use.

One aliquot of each library was thawed, inoculated in SRaf –trp – ura medium and grown for 32h at 28 °C. Afterwards, expression was induced by diluting the cultures to an OD600 of 0.67/mL into SRaf/Gal –trp –ura medium and grown for 16h at 28 °C. The cells were harvested and washed thrice with FACS washing buffer (1X PBS + 1 mM EDTA, pH 7.2 + 1 Complete Inhibitor EDTA-free tablet (Roche) per 50 mL buffer). Then, the cells were incubated at an OD_600_ of 8/mL in FACS staining buffer (washing buffer + 0.5 mg/mL of Bovine Serum Albumin) with 9.09 nM hACE2-muFc (Sino Biological) for 1h. Cells were washed thrice with FACS staining buffer and incubated with 1:100 anti-cmyc-FITC (Immunology Consultants Lab), 1:1000 anti-mouse-IgG-AF568 (Molecular Probes) and 1:2000 L/D eFluor506 (Thermo Fischer Scientific) in staining buffer for 1h. After washing thrice with staining buffer, cells were sorted on a FACSMelody (BD Biosciences), with a selection gate capturing the ACE2+ cells, such that, after compensation, max 0.1% of cells of unstained and single stained controls appeared above the background. Approximately 1 million ACE2+ cells were isolated per library, which were recovered in liquid SD –trp –ura drop-out medium supplemented with 100 U/mL penicillin and 100 μg/mL streptomycin (Thermo Fisher Scientific) for 36h at 28 °C. Afterwards, the cells were snap frozen at 5e7 cells per aliquot and stored at -80 °C for further use.

Similarly as written above, one aliquot of each ACE2+ presorted library was thawed, inoculated and induced. After washing, the cells were incubated at an OD600 of 8/mL in FACS staining buffer with the VHH-Fc molecule for 1h. Cells were washed thrice with FACS staining buffer and incubated with 1:100 anti-cmyc-FITC (Immunology Consultants Lab), 1:1000 anti-human-AF594 (Molecular Probes) and 1:2000 L/D eFluor506 (Thermo Fischer Scientific) in staining buffer for 1h. After washing thrice with staining buffer, cells were sorted on a FACSMelody (BD Biosciences), with a selection gate capturing the ‘escapers’, chosen such that, after compensation, max. 0.1% of cells of the fully stained WT RBD control appeared in the selection gate. Approximately 80.000 – 370.000 cells were captured per library, which were recovered in liquid SD –trp –ura drop-out medium supplemented with 100 U/mL penicillin and 100 μg/mL streptomycin (Thermo Fisher Scientific) for 16h at 28 °C before plasmid extraction.

Plasmid was extracted using the Zymoprep yeast plasmid miniprep II kit (Zymo Research) according to the manufacturer’s instructions, but with a longer (2h) incubation with the Zymolase enzyme, and with the addition of a freeze-thaw cycle in liquid nitrogen after Zymolase incubation. A PCR with the KAPA HiFi HotStart ReadyMix using NEBNext UDI primers (20 cycles) was performed to isolate the barcode and add sample indices and remaining Illumina adaptor sequences. Clean-up was performed with 1.8X CleanNA beads. Hundred bp single-end sequencing was performed on an Illumina NovaSeq 6000 by the VIB Nucleomics Core.

### Analysis of deep mutational scanning to estimate VHH-Fc epitopes

Deep sequencing reads were processed as described by Greaney *et al*.^21^ using the code available at https://github.com/jbloomlab/SARS-CoV-2-RBD_MAP_Crowe_antibodies, with adjustments. Briefly, nucleotide barcodes and their corresponding mutations were counted using python (3.7.6) and dms_variants package (0.8.6). Escape fraction for each barcode was defined as the fraction of reads after enrichment divided by the fraction of reads before enrichment of escape variants. The resulting variants were filtered to remove unreliably low counts and keep variants with sufficient RBD expression and ACE2 binding (based on published data^12^). For variants with several mutations, the effects of individual mutations were estimated with global epistasis models, excluding mutations not observed in at least one single mutant variant and two variants overall. The resulting escape measurements correlated well between the duplicate experiments and the average across libraries was thus used for further analysis. To determine the most prominent escape sites for each VHH-Fc, RBD positions were identified where the total site escape was > 10x the median across all sites, and was also at least 10% of the maximum total site escape across all positions for a given VHH-Fc.

### Hamster challenge

The hamster infection model of SARS-CoV-2 as described before ^22^. In brief, female Syrian hamsters (*Mesocricetus auratus*) of 6-8 weeks old were anesthetized with ketamine/xylazine/atropine and inoculated intranasally with 50 μL containing 2×10^6^ TCID50 SARS-CoV-2 BetaCov/Belgium/GHB-03021/2020. Animals were treated once by intraperitoneal injection, either 24h post SARS-CoV-2 challenge. Hamsters were monitored daily for appearance, behavior and weight. At day 4 post-infection, hamsters were euthanized by intraperitoneal injection of 500 μL Dolethal (200mg/ml sodium pentobarbital, Vétoquinol SA).

Lung samples were collected, and viral RNA and infectious virus were quantified by RT-qPCR and end-point virus titration, respectively. RNA was isolated (nucleic acid purification on the MagNA Pure 96; Roche Life Science), reverse transcribed and Taqman PCR (on the 7500 RealTime PCR system; Applied Biosystems) was performed using specific primers (E_Sarbeco_F: 5′ACAGGTACGTTAATAGTTAATAGCGT3′ and E_Sarbeco_R: 5′ATATTGCAGCAGTACGCACACA3′) and a probe (E_Sarbeco_P1: 5′ACACTAGCCATCCTTACTGCGCTTCG3′) specific for betacoronavirus E gene. Quadruplicate 10-fold serial dilutions were used to determine the virus titers in confluent layers of Vero E6 cells. To this end, serial dilutions of the samples were made and incubated on Vero E6 monolayers for 1hat 37 °C. Vero E6 monolayers were then washed and incubated for 4-6 days at 37 °C after which plates were stained and scored using the vitality marker WST8 (colorimetric readout). To this end, WST-8 stock solution was prepared and added to the plates. Per well, 20 μl of this solution (containing 4 μl of the ready-to-use WST-8 solution from the kit and 16 μl infection medium, 1:5 dilution) was added and incubated 3-5 hours at RT. Subsequently, plates were measured for optical density at 450 nm (OD_450_) using a microplate reader and visual results of the positive controls (cytopathic effect (cpe)) were used to set the limits of the WST-8 staining (OD value associated with cpe). Viral titers (TCID50/ml or/g) were calculated using the method of Spearman-Karber ^23^.

## Competing interests

C.L., H.E., K.R., B.S., S.D.C., D.F., W.N., J.N., X.S., L.v.S., and N.C. are named as inventors on patent application Coronavirus Binders (WO 2021/156490 A2), published on 12 August 2021. C.L., B.S., W.W., L.v.S. and N.C. are named as inventors on patent application Engineered stabilizing aglycosylated fc-regions (WO 2023/148397 A1), published on 10 August 2023.

X.S. and N.C. are scientific founders of and consultants for ExeVir Bio and are in receipt of ExeVir Bio share options.

All other authors declare that they have no competing interests.

## Acknowledgments

We are grateful to Jesse Bloom for providing the *S. cerevisiae* RBD display library and to Arne Matthys, Justine Naessens and Charlotte Roels for providing nanobody sequences. We thank Caroline Foo, Carolien De Keyzer, Lindsey Bervoets, Birgit Voeten, Thibault Francken and Lotte Coelmont for technical assistance with the hamster challenge studies.

We thank the VIB Nanobody Core, the staff of the VIB Flow Core Ghent for providing access to flow cytometry equipment and for their technical assistance, the staff of the VIB Nucleomics Core for Illumina sequencing, and the VIB Protein Service Facility and Savvas Savvides for making MALS and BLI equipment available.

## Funding

This project was supported by the Bill and Melinda Gates Foundation (INV-037592), by the EOS joint programme of Fonds de la recherche scientifique—FNRS and Fonds wetenschapellijk onderzoek–Vlaanderen—FWO (G0H7518N EOS ID: 30981113 and G0H7322N, EOS ID: 40007527) to X.S., by FWO project G0B1917N to X.S.; by FWO project G0G4920N to X.S., N.C. and J.N.

C.L. was supported by a Strategic Basic Research fellowship of the Fund for Scientific Research Flanders (FWO) (application number 1S11418N) and the Gates foundation. H.E. was supported by a Fundamental fellowship of the FWO (11C9720N). S.D.C and I.R. were supported by a PhD fellowship of the FWO.

