## Supplementary Figures for "Yeast-based production platform for potent and stable heavy chain-only antibodies"

```
XVR011       1    DVQLVESGGGLVQPGGSLRLSCAASGRTFSEYAMGWFRQAPGKEREFVATISWSGGATYY    60

XVR011       121  VTVSS--GGGGSGGGGSDKTHTCPPCPAPEAAGGPSVFLFPPKPKDTLMISRTPEVTCVVVD  180

VHH1-FcGen1  121  .....GS.......................LL..............................  182
VHH1-FcGen2  121  .....GS.......................LL..............................  182

XVR011       181  VSHEDPEVKFNWYVDGVEVHNAKTKPREEQYNSTYRVVSVLTVLHQDWLNGKEYKCKVSN    240

VHH1-FcGen1  183  ...............................A............................    242
VHH1-FcGen2  183  ..........................C....A....C.......................    242

XVR011       241  KALPAPIEKTISKAKGQPREPQVYTLPPSRDELTKNQVSLTCLVKGFYPSDIAVEWESNG    300

VHH1-FcGen1  243  ............................................................    302
VHH1-FcGen2  243  ............................................................    302

XVR011       301  QPENNYKTTPPVLDSDGSFFLYSKLTVDKSRWQQGNVFSCSVMHEALHNHYTQKSLSLSPG   361

VHH1-FcGen1  303  .............................................................K  364
VHH1-FcGen2  303  .............................................................K  364
```

**Supplementary Figure S1. Alignment of the amino acid sequences of the Fc-engineered anti-Covid VHH-Fc molecule to XVR011.**

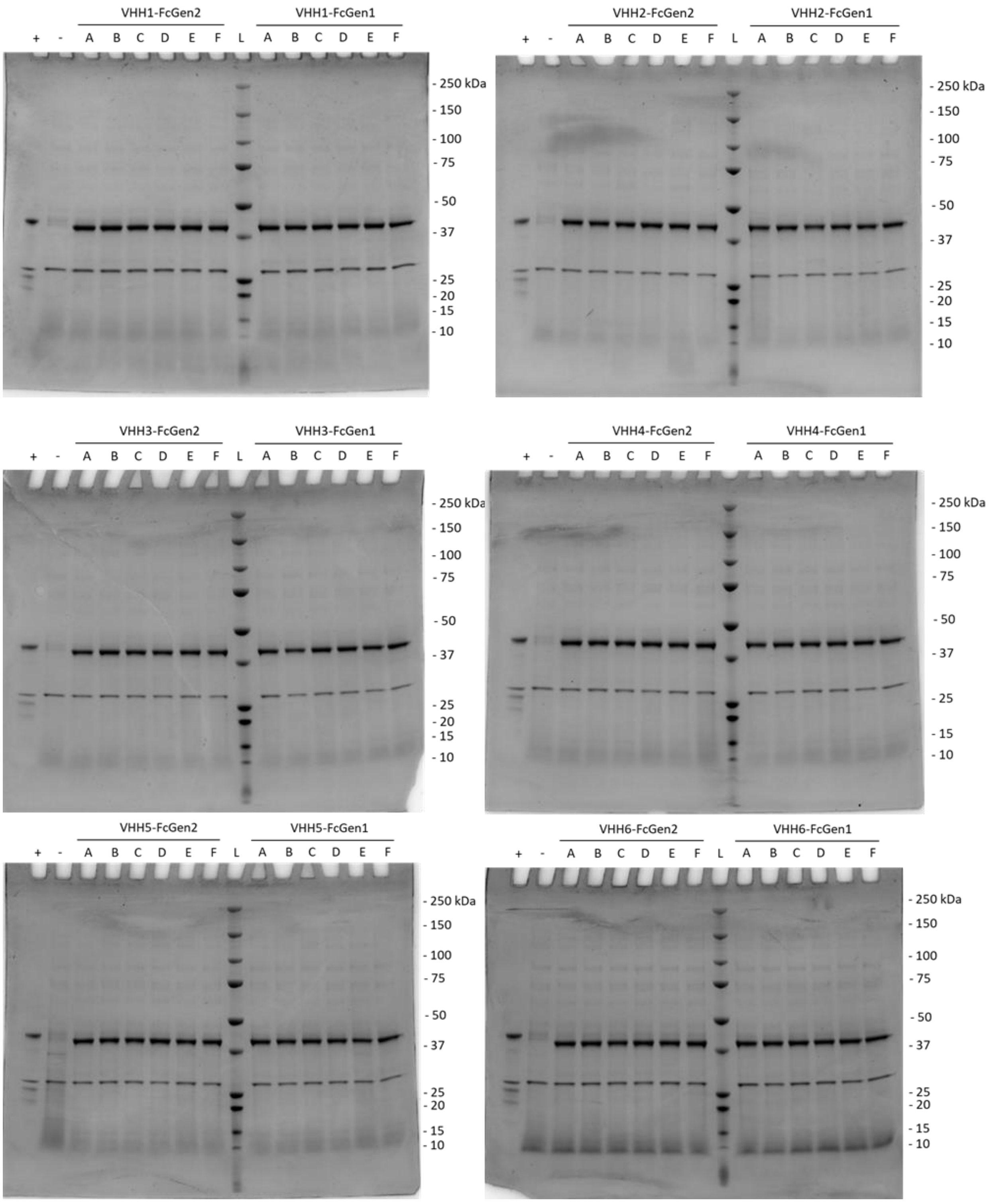

**Supplementary Figure S2. Reducing SDS-PAGE of supernatant of randomly selected clones of *Pichia pastoris* strain *OPENPichia* expressing VHH-Fc clones (A-F) expressed under the $P_{GAP}$ promotor.** EndoH (0.5 µg) was used as a loading control. Ladder is Precision Plus Protein™ All Blue Prestained Protein Standards. Negative control (-) is wild-type *Pichia* supernatant. Positive control (+) is a purified VHH-Fc molecule.

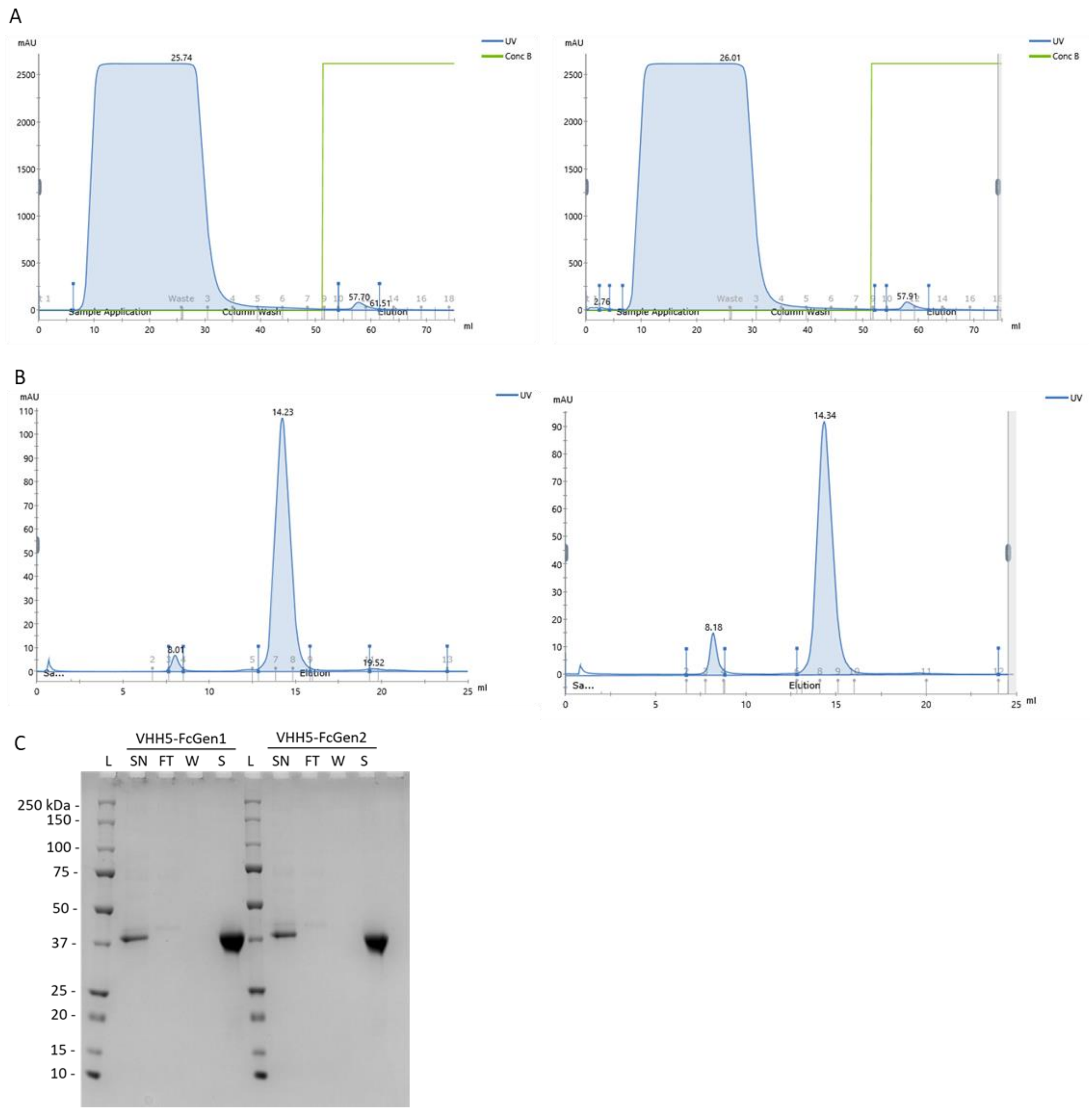

**Supplementary Figure S3. Purification of VHH-Fc molecules.**

A. UV-chromatograms of the supernatant of *Pichia pastoris* (strain OPENPichia) containing VHH-Fcs loaded on a protein A column. Example (VHH5-FcGen1 and VHH5-FcGen2) is representative for all produced VHH-Fcs.

B. UV-chromatograms of protein A purified VHH-Fcs loaded on a size exclusion chromatography resin. Example (VHH5-FcGen1 and VHH5-FcGen2) is representative for all produced VHH-Fcs.

C. Reducing SDS-PAGE of samples taken during purification. Example (VHH5-FcGen1 and VHH5-FcGen2) is representative for all produced VHH-Fcs. SN is supernatant, FT is flow-through, W is protein A wash and S is SEC peak fraction. Ladder (L) is Precision Plus Protein™ All Blue Prestained Protein Standards.

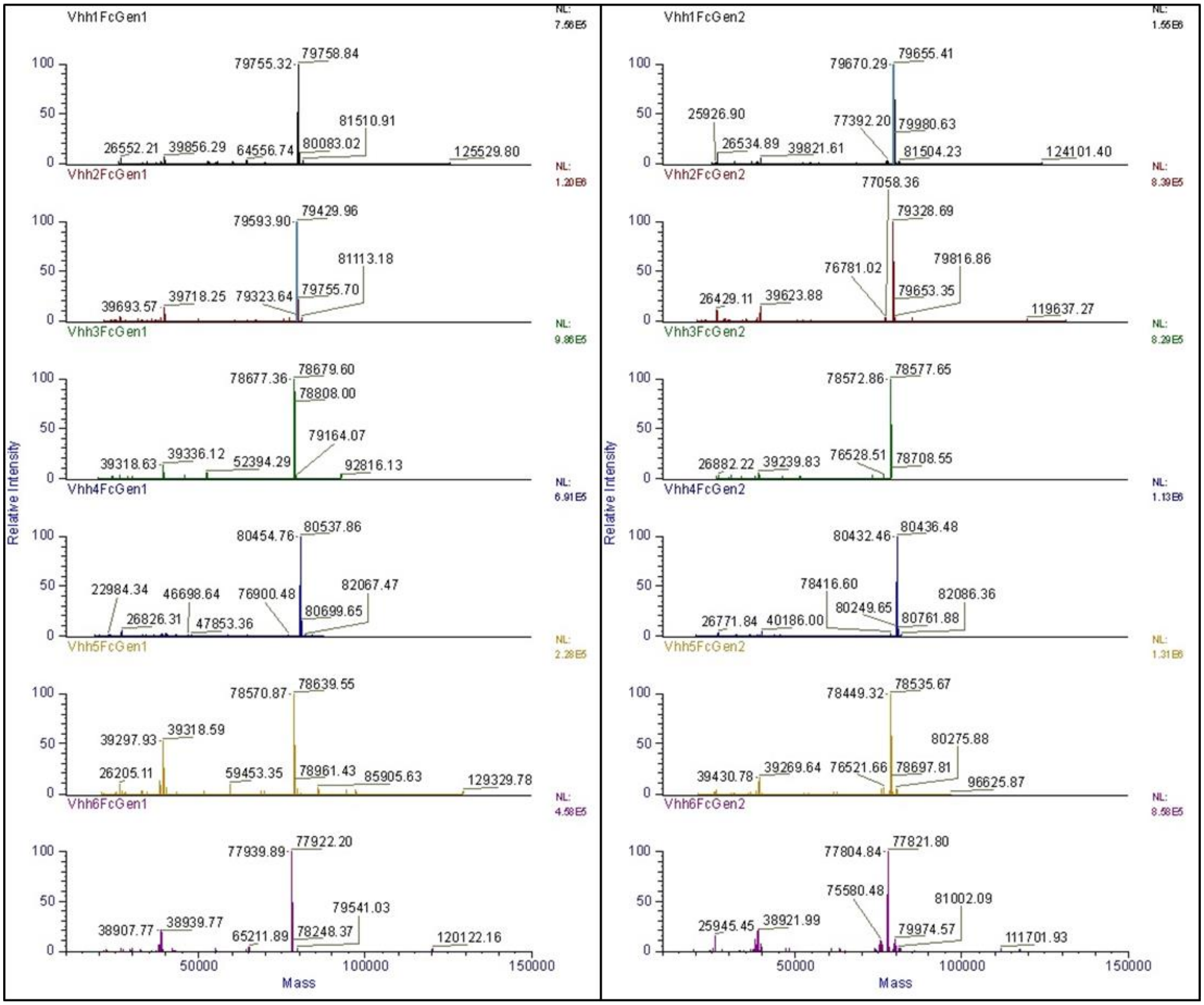

**Supplementary Figure S4. LC-MS spectra of intact VHH-Fcs produced by *Pichia pastoris* strain OPENPichia and purified via protein A and size exclusion chromatography.** The raw spectra were deconvoluted with the ReSpect Algorithm in BioPharma Finder software, followed by manual annotation.
